# The mitotic crosslinking protein PRC1 acts as a mechanical dashpot to resist microtubule sliding

**DOI:** 10.1101/2019.12.12.874529

**Authors:** Ignas Gaska, Mason Armstrong, April Alfieri, Scott Forth

## Abstract

Cell division in eukaryotes requires the regulated assembly of the spindle apparatus. The proper organization of microtubules within the spindle is driven by motor proteins that exert forces to push and slide filaments, while non-motor proteins can crosslink filaments into higher order motifs such as overlapping bundles. It has not been clear how active and passive forces are integrated to produce regulated mechanical outputs within spindles. Here we employ a combined optical tweezers and TIRF microscopy instrument to directly measure the resistive forces produced by the mitotic crosslinking protein PRC1. We observe that PRC1 generates frictional forces that resist microtubule sliding. These forces scale with microtubule sliding velocity and the number of PRC1 crosslinks, but do not depend on overlap length or PRC1 density within overlaps. Our results suggest that PRC1 ensembles act like a mechanical dashpot, producing significant resistance against fast motions, but minimal resistance against slow motions, allowing for the integration of diverse motor activities into a single mechanical outcome.

## Introduction

Cell division in eukaryotes requires the organization of the microtubule-based mitotic spindle to segregate chromosomes and regulate positioning of the cell division plane (Kapoor, 2017). Within the spindle, subsets of microtubules are organized into specialized arrays whose filaments can undergo relative sliding (McIntosh and Hays, 2016). Microtubule sliding is driven in part by motor proteins that either crosslink filaments or cluster at microtubule ends to exert active sliding or pulling forces along the filament (Forth and Kapoor, 2017). Single molecule analyses of diverse motor proteins found within spindles indicate that stepping rates can vary over nearly several orders of magnitude, from dynein at nearly ∼1 micron/sec (King and Schroer, 2000; McKenney et al., 2014; Ross et al., 2006) to various mitotic kinesins that range from 25-300 nm/sec (Braun et al., 2017; Drechsler et al., 2014; Kapitein et al., 2005; Reinemann et al., 2017, 2018; Wijeratne and Subramanian, 2018). Spindle microtubules can also be bundled into higher-order arrays by non-motor proteins that can sort filaments by polarity and regulate the relative motions of sliding microtubules (Subramanian and Kapoor, 2012). It has been proposed that crosslinking by such proteins may produce a ‘brake-like’ resistance to filament sliding that acts to balance motor-driven forces (Janson et al., 2007), leading to an integration of active and passive forces that sets parameters such as spindle length and filament sliding velocity.

Among the non-motor mitotic crosslinking proteins are members of the conserved MAP65 family, which include the yeast Ase1 and the human PRC1. These proteins have been shown to preferentially crosslink antiparallel microtubules both in vitro and within the anaphase central spindle microtubule array (Bieling et al., 2010; Janson et al., 2007; Rincon et al., 2017; Subramanian et al., 2010; Tikhonenko et al., 2016; Yamashita et al., 2005) and to help form “bridging fibers” to crosslink kinetochore microtubules during metaphase (Kajtez et al., 2016; Polak et al., 2017; Vukušić et al., 2017). Several lines of evidence suggest that microtubule arrays crosslinked by MAP65 proteins act as mechanical linkages between segregating chromosomes and separating spindle halves in anaphase. First, knockdown of PRC1 in human cell lines reveals that microtubule density is reduced within spindle midzones in anaphase and the two spindle halves appear fully detached from one another (Mollinari, 2004; Mollinari et al., 2002; Zhu et al., 2006) and chromosome segregation rates and separation distance both increase (Pamula et al., 2019). Second, the yeast MAP65, Ase1, has been shown in in vitro assays to slow motor-driven microtubule sliding via an adaptive braking mechanism (Braun et al., 2011) and can autonomously produce entropic expansion forces that prevent microtubules from sliding apart (Lansky et al., 2015). In dividing yeast cells, Ase1 has been shown to selectively stabilize longer bundles while still allowing for the motor-driven transport of shorter microtubules, suggesting that Ase1 can produce an overlap length-dependent resistance (Janson et al., 2007). Third, single molecule studies revealed that monomeric PRC1 constructs containing just the microtubule-binding domain of the protein generate frictional resistance when moved along the microtubule lattice when under tension (Forth et al., 2014). However, it is currently unclear how ensembles of full-length human PRC1 molecules crosslinking microtubules in micron-scale overlaps produce resistive forces. Specifically, it is unknown how resistive forces might be regulated within sliding bundles by parameters such as PRC1 density, microtubule overlap length, or relative sliding speed.

Here, using simultaneous TIRF microscopy and optical trapping methods, we directly measure the resistive forces generated by ensembles of PRC1 molecules crosslinking microtubule pairs. We find that the magnitude of resistive forces linearly scales both with the velocity at which microtubules slide apart and with the total number of engaged crosslinkers in overlap region, but not the length of the overlap or the density of crosslinks. Surprisingly, we observe that pausing during sliding allows the system to relieve tension and reinitiating sliding results in increased resistive forces, suggesting that upon cessation of relative sliding, PRC1 molecules can rearrange within the overlap to form a higher-order configuration that more strongly resists microtubule separation. Finally, using computational modeling, we describe how PRC1 can be condensed into shrinking overlaps by microtubule ends that act as ‘leaky’ reflective barriers to diffusion. Together, these results suggest that PRC1-crosslinked microtubule pairs generate viscous resistance to act like a mechanical dashpot during filament sliding.

## Results

### Relative sliding of microtubules crosslinked by PRC1 generates resistive forces

To characterize the mechanical properties of PRC1-mediated microtubule bundles, we established an assay that allowed us to control microtubule sliding motions while simultaneously measuring the forces between two crosslinked microtubules and observing the distribution of GFP-PRC1 molecules within microtubule overlap regions. To achieve this, we employed a strategy similar to one previously used to measure pushing forces by kinesin-5 ensembles (Shimamoto et al., 2015). Briefly, microtubules containing HiLyte-647 and biotinylated tubulin were immobilized on a passivated coverslip via Neutravidin linkages. GFP-PRC1 and rhodamine-labelled microtubules were introduced into the sample chamber, generating microtubule bundles with varying overlap lengths and amounts of PRC1 molecules in overlaps. Finally, polystyrene beads coated with truncated kinesin-1 construct were introduced, allowing for long-lived attachments to microtubules. These beads were then held in an optical trap and used to both apply and measure force along the length of the microtubule bundle (Figure 1A).

**Figure 1.**
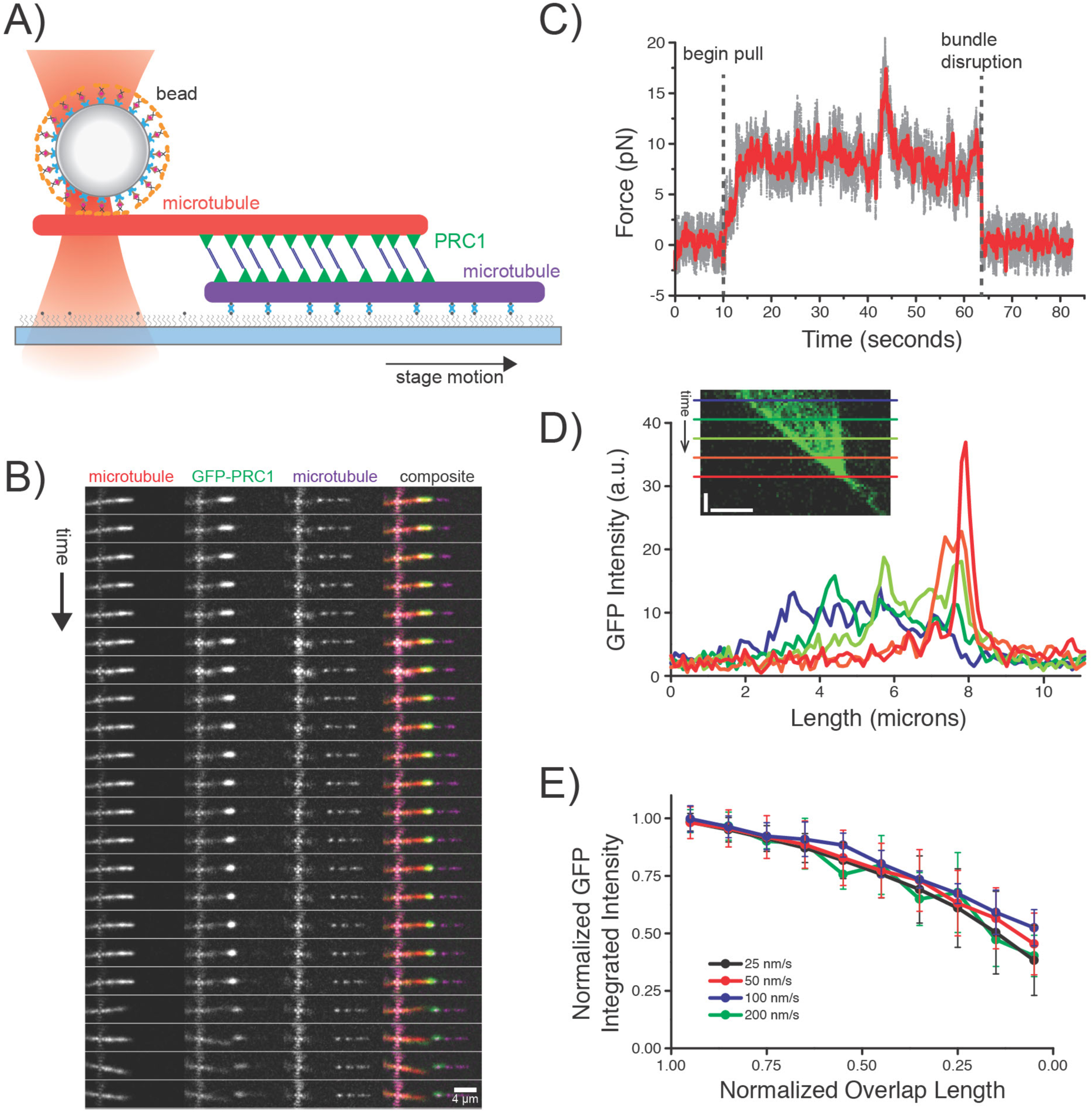
Direct measurement of force across sliding microtubules crosslinked by PRC1. (A) Schematic of the assay. A surface-immobilized microtubule (labelled with HiLyte-647 tubulin and biotinylated-tubulin, purple) is crosslinked via GFP-PRC1 molecules to a second microtubule (labelled with rhodamine-tubulin, red) which is held via a kinesin-coated bead in an optical trap. (B) Time series of fluorescent images acquired via TIRF microscopy during microtubule sliding event. GFP-PRC1 and individual microtubule channels are shown along with composite image. Scale bar = 4 microns. (C) Representative force time series data (raw, gray; averaged, red) during microtubule sliding. Initiation of pull at 10s and disruption event at ∼63s are labeled. (D) Linescan analysis from GFP channel. GFP intensities at select time points are plotted against length, with corresponding time points noted on kymograph inset (blue = 5s, green = 15s, yellow = 25s, orange = 35s, red = 45s). Kymograph scale bars: vertical = 10s, horizontal = 2 microns. (E) GFP signals within overlap regions were integrated and normalized to their values at the initial overlap and are plotted against normalized overlap length. Data from sliding events at four different velocities are shown (N>8 for each velocity, error bars = SD).

Moving the sample stage at a fixed velocity parallel to the microtubule bundle axis allowed for controlled separation of filaments. Images of each of the two microtubules and the GFP-PRC1 molecules were acquired using total internal reflection fluorescence (TIRF) microscopy. These data show the surface-immobilized microtubule moving at a constant velocity, the free microtubule held in place via the optically trapped bead, and the GFP-PRC1 molecules clustering into the region of decreasing overlap (Figure 1B). Upon complete separation of a microtubule pair, the free microtubule became detached from the bundle and freely swiveled, while a small number of remaining GFP-PRC1 molecules bound to the surface microtubule began to diffuse away from the microtubule end. Simultaneously with the acquisition of the fluorescence imaging data, we recorded the force exerted on the bead using a custom-built force-calibrated optical tweezers instrument (Figure 1C). Before separating the microtubule pair, the measured force was approximately 0 pN. After a 10-second delay to confirm bead attachment and record the initial distribution of microtubules and PRC1 molecules, the surface microtubule was moved at a constant velocity and the magnitude of the force increased rapidly until reaching a relatively stable value. Over the course of the pulling event, the force remained nearly constant, with small and stochastic deviations around an average value. Upon bundle separation, the force value sharply dropped to 0 pN, consistent with the microtubules no longer being mechanically crosslinked. Together, these data demonstrate that our instrument is capable of simultaneous force and fluorescence measurement of sliding microtubule bundles moving at controllable velocities, and that crosslinked microtubule pairs generate resistive forces when sliding.

We next examined the time-dependent distribution of GFP-PRC1 molecules within the overlap region throughout the experiment. Line scans of the GFP signal intensity were generated for each time point and plotted against distance along the microtubule positions (Figure 1D and kymograph inset). We observed that at earlier time points, when the overlap region is several microns in length, the GFP intensity is relatively evenly distributed within the overlap and has a mean value of ∼10 a.u. (blue line). As the overlap decreased in length, the mean value of the intensity increased while the length distribution became narrower, indicating that PRC1 molecules track the shrinking overlap region. Once the overlap had reached a value close to 0 microns, the GFP intensity signal reached a maximum value of ∼40 a.u. (red line). Control experiments with non-moving microtubule pairs reveal that loss of GFP fluorescence signal due to photobleaching occurs at a rate of only ∼4% per minute, suggesting that time-dependent changes in signal are due to changes in PRC1 concentration, and not photobleaching (Figure S1).These data suggest that the PRC1 molecules are moving closer together within the overlap, increasing in density as the overlap length decreases.

We next sought to determine what percentage of PRC1 molecules persisted within the overlap during its reduction in length. We calculated the integrated GFP intensity within overlaps, normalized both the integrated intensity and overlap lengths relative to their initial values, and compared these relationships at four different pulling velocities (Figure 1E). Here, we observe that as the overlap length decreases, the percentage of remaining PRC1 molecules decreases. As the overlap length approached zero, approximately one third to one half of the original molecules can still be detected within the overlap. Together, these data suggest that a subset of the initial population of PRC1 molecules is lost from the bundle, but the molecules that remain are driven to cluster within the shrinking overlap region between sliding microtubules.

### The magnitude of resistive forces increases with increasing relative microtubule velocity

We next sought to determine how resistive forces generated during bundle disruption would depend on sliding velocity. We therefore disrupted microtubule bundles at four different velocities that are similar to reported stepping rates for various mitotic kinesins: 25 nm/s, 50 nm/s, 100 nm/s, and 200 nm/s. Inspection of the fluorescence data revealed that at each velocity, bundle overlap decreased uniformly at the rate of microtubule sliding, and PRC1 molecules consistently tracked and concentrated within the shrinking overlap (Figure 2A). Forces measured during bundle disruption events appeared to be generally larger in magnitude at higher sliding velocities (Figure 2B). We confirmed that the bead-microtubule attachments persisted throughout all pulls by noting that the overlap length reduction was continuous and no sudden force reductions to zero load were observed, as we saw in control cases where bead-microtubule attachment was transiently lost (Figure S2). We also confirmed that the microtubule-surface attachments remained robust throughout our experiments by measuring the velocity of the surface-immobilized microtubule and finding it to consistently match that of the controlled stage velocity (Figure S3).

**Figure 2.**
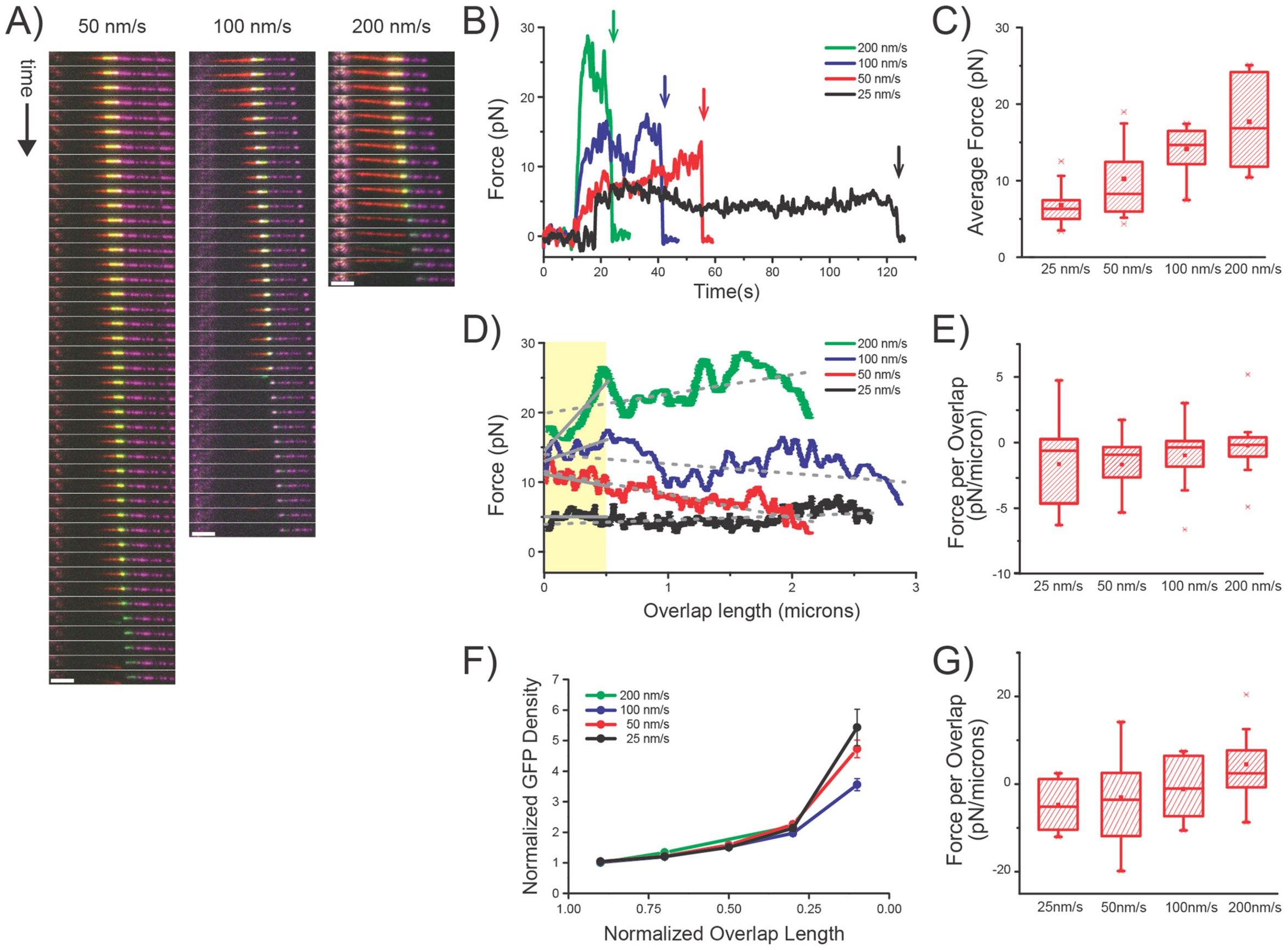
Resistive forces generated by PRC1 crosslinks scale with relative microtubule sliding velocity. (A) Time series of composite fluorescent images acquired via TIRF microscopy during microtubule sliding events at three different velocities: 50 nm/s, 100 nm/s, and 200 nm/s. Purple = surface-immobilized microtubule, red = bead-bound microtubule, green = GFP-PRC1. (B) Representative averaged force time series at four velocities (black = 25, red = 50, blue = 100, green = 200 nm/s). Disruption events are marked with vertical arrows. (C) Box plots of mean force values during sliding events plotted for each velocity (N>7 for all velocities). (D) Representative force trajectories plotted against measured bundle overlap length at four different velocities, (black = 25, red = 50, blue = 100, green = 200 nm/s). Shaded yellow region highlights bundle overlap lengths of 0-0.5 microns. Dashed gray lines = linear fits for duration of entire bundle pull event. Solid gray lines = linear fits for bundle overlap lengths of 0-0.5 microns. (E) Box plots of slopes calculated from force versus total overlap length (N>9 for all velocities). (F) GFP Density values normalized to their values at the initial overlap are plotted against normalized overlap length. Data from all events were binned in 200nm length increments and averaged for each velocity. (N>7 for all velocities, error bars = SE). (G) Box plots of slopes calculated from force versus overlap lengths for data selected between 0-0.5 microns of overlap length (N>7 for all velocities).

We repeated these experiments multiple times for each condition and calculated the average force during sliding events at each velocity (Figure 2C), revealing a positive correlation between faster sliding velocity and increasing force. We next considered the relationship between force and overlap length for the same data sets. These data reveal that the force trend throughout the entire sliding event remains relatively constant, suggesting minimal correlation between force and overlap length (Figure 2D). After calculating the slopes of these data (Figure 2D, dashed gray lines) and averaging multiple traces, we confirmed that the change in force with overlap length is close to 0 pN/micron, but with a slightly negative trend (Figure 2E). This indicates that there may be a small increase in force as the bundle approaches separation, but that the force is relatively constant within a given pulling event and is largely independent of overlap length. Previous analyses of microtubule bundles crosslinked by Ase1 molecules suggested that as the density of crosslinkers increased, entropic forces that resisted bundle separation rapidly grew in magnitude (Lansky et al., 2015). We observe a similar increase in the density of PRC1 molecules as overlap lengths approach zero (Figure 2F). However, analysis of the change in force values relative to change in overlap lengths for data subsets selected between 0 and 0.5 microns (Figure 2D, solid grey lines) do not reveal a sharp increase as overlap lengths approach 0 microns (Figure 2G), suggesting that the resistance to microtubule pair separation produced by Ase1 and PRC1 operate by different mechanisms.

### Resistive forces scale with number of PRC1 molecules but not overlap length

While we observed a general trend of force values that increase with faster pulling velocities, we noted that there were individual instances where the force recorded during pulls at our slowest rate (25 nm/s) exhibited higher average values than those measured during our fastest pulls (200 nm/s) (Figure 2C). This led us to examine additional parameters that may contribute to the production of resistive forces. The formation of bundles occurs spontaneously within the sample chamber, and therefore bundles can have both variable initial overlap lengths and different concentrations of GFP-PRC1. To allow for more direct comparisons across different pulling rates, we selected subsets from our data to perform analyses within a small range of overlap values (0.5-1.5 microns). We next plotted individual force and GFP integrated intensity data points from within this range for each velocity and found that forces increased with increasing integrated GFP intensity (Figure 3A). We next calculated the slopes of each constant sliding velocity data set and observed that the average slope value increased nearly linearly with increasing velocity (Figure 3B). Plotting the same force values against overlap length did not show a significant correlation (Figure 3C), and the slopes of the force-overlap relationships within this subset of data did not show a clear dependence on overlap length (Figure 3D). Together, these data reveal that the magnitude of resistive force scales both with GFP integrated intensity, which is proportional to the total number of PRC1 molecules in the overlap, as well as sliding velocity, but does not depend on the overlap geometry of the system.

**Figure 3.**
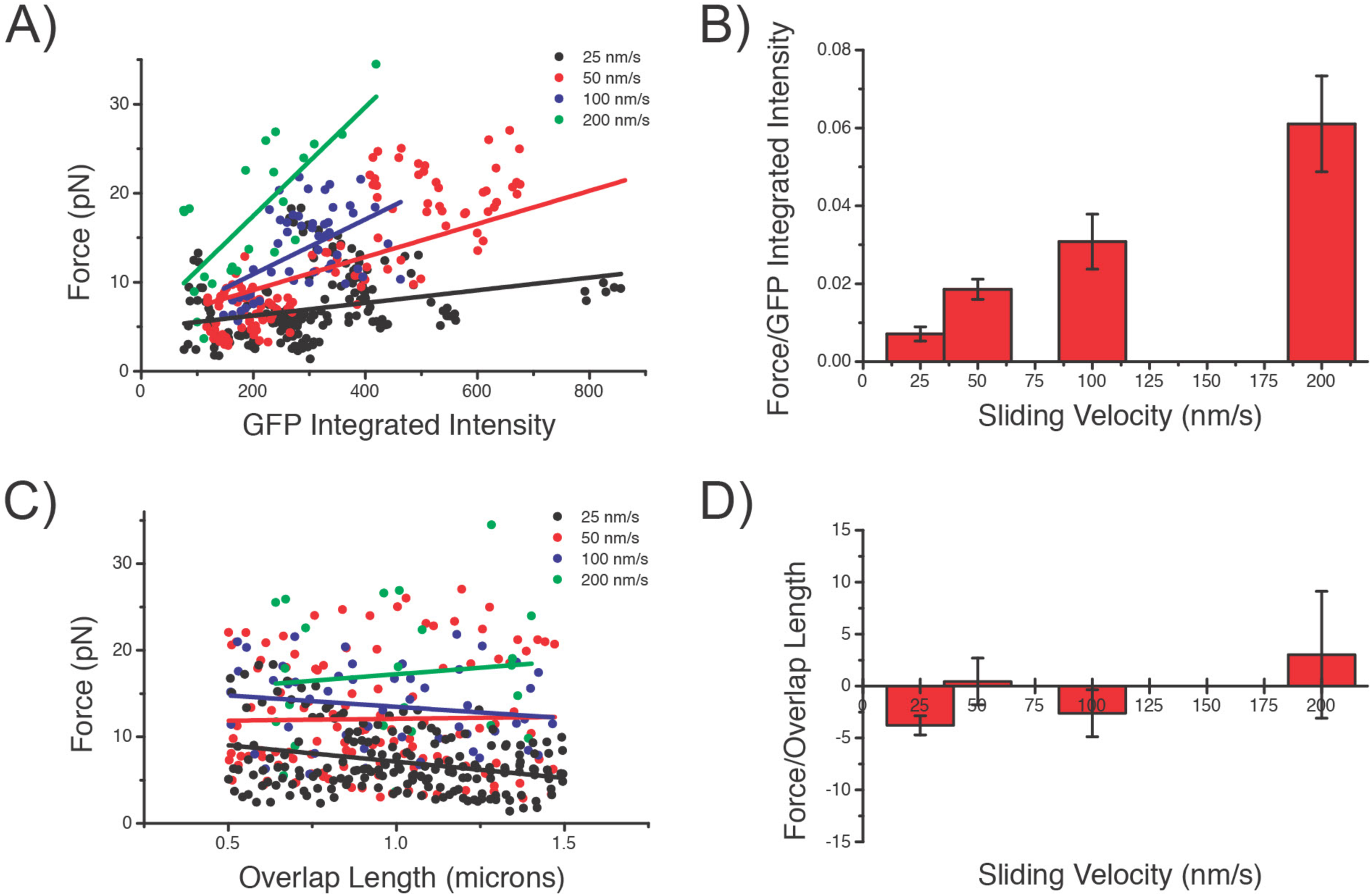
Resistive forces are proportional to PRC1 crosslink number but not overlap length. (A) Integrated GFP intensity values within overlaps and the corresponding average force during image acquisition period. Data from overlaps between 0.5 and 1.5 microns from all traces are plotted (Points: black = 25 nm/s, red = 50 nm/s, blue = 100 nm/s, green = 200 nm/s. Solid lines = linear fits to each data set). (B) Slopes of linear fits from (A). (C) Same averaged force data points from (A), plotted against corresponding overlap length (Points: black = 25 nm/s, red = 50 nm/s, blue = 100 nm/s, green = 200 nm/s. Solid lines = linear fits to each data set). (D) Slopes of linear fits from (C). N>8 for all velocities.

### Pausing of sliding results in force relaxation, while resumption of sliding produces increased resistive forces

We occasionally encountered situations where attempts to slide bundles apart failed, due to insufficient force production from our trapping laser or bead detachment from the microtubule due to high loads generated despite no observable filament sliding. These events became more frequent the longer the sample sat on the microscope, leading us to hypothesize that PRC1 molecules within overlaps were undergoing a time-dependent rearrangement that was more resistant to microtubule sliding. To test this idea, we repeated our sliding experiments but introduced a short (20 second) pause in the microtubule motion before resuming the sliding event. Measured forces revealed three distinct regimes: an initial sliding event similar to what we’d previously observed, a relaxation event during the pause, and a second sliding event whose force magnitude and rate of force increase were larger than for the initial pull (Figure 4A). Fluorescence images reveal that the overlap region did not appreciably change in length, nor did the integrated GFP signal decrease during the pause (Figure 4B). These distinct behaviors between the first and second pull were consistent across all bundles tested (Figure 4C). During the pause, we observed a reduction in force that decayed to nearly zero pN after ∼20 seconds (Figure 4D). Normalizing the force values relative to the initial value at the pause onset revealed a characteristic exponential decay whose time constant was similar (7.5 +/−0.2 seconds) for all bundles examined, regardless of the magnitude of the initial force (Figure 4E). Finally, we quantified the mean force measured with each of the constant sliding regimes (Figure 4F) and the rate of force increase as measured by the slope of the force signal prior to reaching a plateau value (Figure 4G). For each of these parameters, the values measured during the second pull were consistently higher relative to the first pull. Calculating the ratio of these values revealed that the mean force and force increase rate were nearly twice as large during the second pull relative to the first (Figure 4H). Together these data suggest that allowing tension within the system to relax after sliding leads to a rearrangement of PRC1 molecules that is capable of producing increased resistive forces upon re-initiation of constant-velocity sliding.

**Figure 4.**
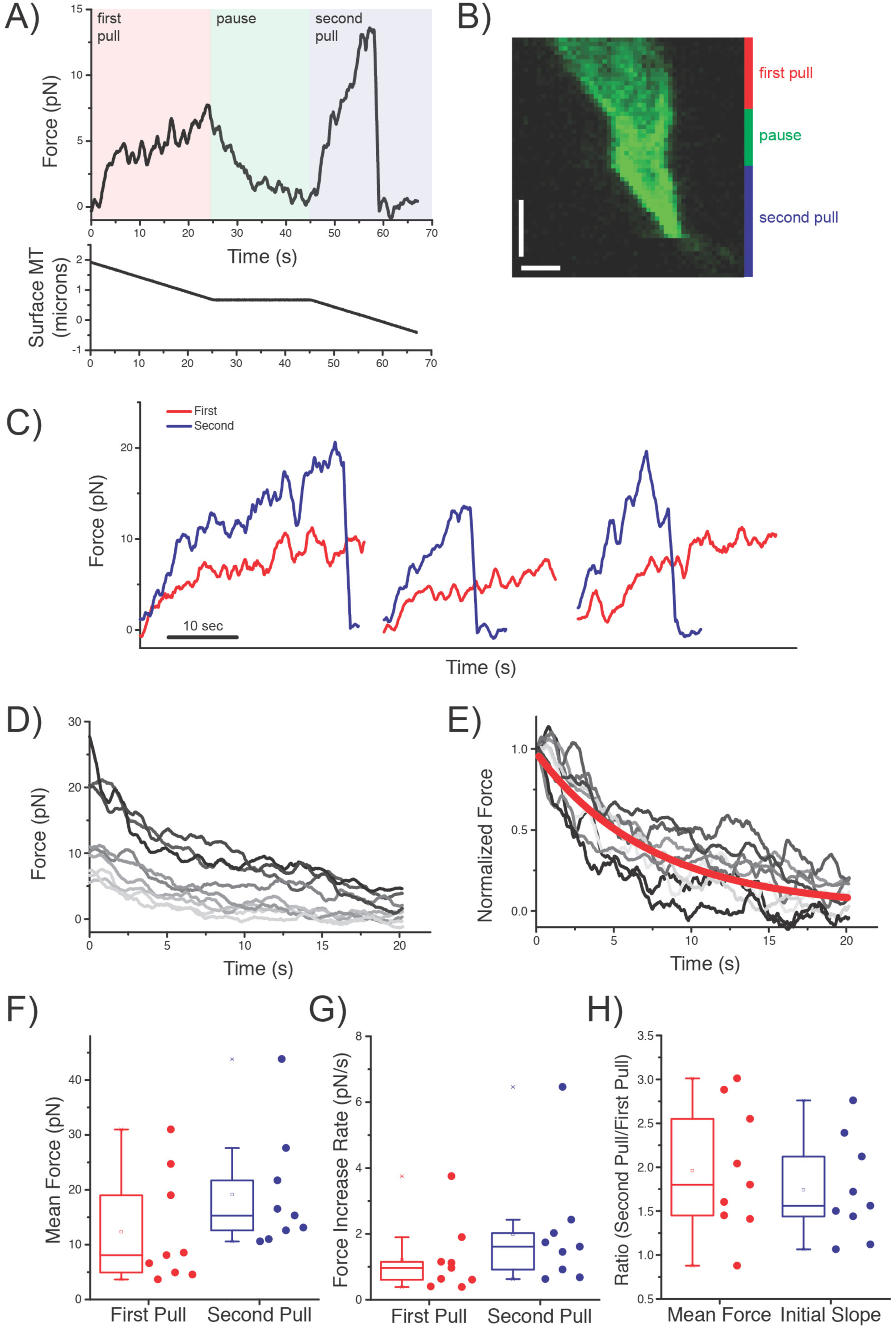
Pausing and then reinitiating sliding results in increased force production after relaxation. (A) Representative force time series for double-pull assay. Shown are force and microtubule positions during initial pull (red shading), 20 second pause (green shading), and second pull (blue shading). (B) Kymograph of GFP fluorescence signal during double pull experiment (vertical scale bar = 20 seconds; horizontal scale bar = 1 micron). (C) Example initial and second pull force time series for three different bundles. Red = initial pull, Blue = second pull. (D) Force time series during pause. Absolute force data are shown (N = 9 independent bundles). (E) Normalized force time series during pause. Red line = exponential decay best fit to all data. Characteristic time = 7.5 +/− 0.2 seconds. (F) Mean force calculated during initial and second sliding events. (G) Rate of force increase calculated from linear fit of force versus time during first 5 seconds of pull for initial and second pulling event. (H) Ratio of values calculated in (F) and (G) for initial and second pulling event (N=9 pull/pause/pull events).

### Computational modeling suggests that a partially reflective end barrier against PRC1 diffusion allows for sustained resistive forces

In order to gain insight into the mechanism by which PRC1 molecules produce resistive forces that scale with velocity but not overlap length, we turned to computational modeling. We expanded upon a previously reported simulation method (Shimamoto et al., 2015) that models two microtubules bundled by individual crosslinkers whose load-dependent behavior on the microtubule lattice can be defined by four parameters. First, an individual crosslinking molecule has two domains which are linked by a linear ‘spring’ whose stiffness is described by k_stiffness_ and which, when separated by a distance Δx, can produce a force F = k_stiffness_*Δx. Second, each of these domains can bind to and diffuse along the microtubule surface with a force-dependent rate k_diffuse_(F). Third, each microtubule-binding domain can detach from the microtubule lattice it contacts with a force-dependent rate constant described by k_detach_(F). Fourth, we introduced a parameter k_end_ which describes the rate at which the molecule can diffuse off of the microtubule end once it reaches the last available lattice site on the microtubule (Figure 5A). Finally, we introduced the constraint that a molecule cannot move into a site that is already occupied by another crosslinking molecule. The simulation could then be run with a variable number of initial crosslinking molecules, overlap length, and rate of relative microtubule sliding.

**Figure 5.**
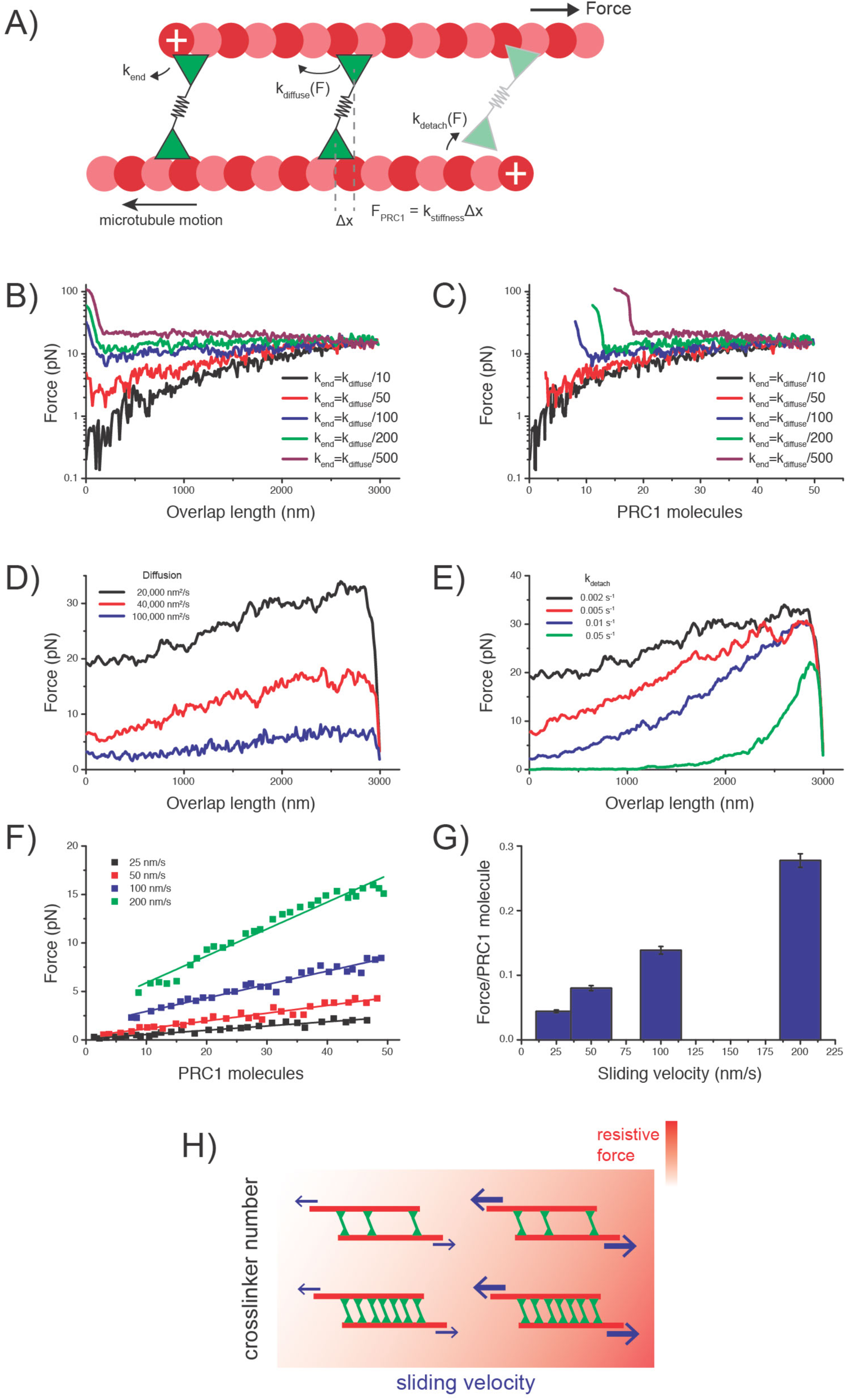
A computational model predicts a partially reflective end barrier against PRC1 diffusion. (A) Schematic depicting parameters used in computational simulations. PRC1 molecules bind to the microtubule lattice on sites with 8 nm periodicity, and can diffuse with rate k_diffuse_(F), detach from the microtubule with rate k_detach_(F), and diffuse off of microtubule ends with rate k_end_. Force across each PRC1 molecule is calculated by F = k_stiffness_ * Δx. (B) Effect of varying k_end_ relative to k_diffuse_. For each parameter set, 20 simulations were run, and values were averaged. Force versus overlap length are shown for a range of k_end_ values. (C) Force data from (B) plotted as a function of engaged PRC1 crosslinking molecules. (D) Effect of varying k_diffuse_ while setting k_detach_ = 0.002 s^−1^. Force versus overlap length are shown for a range of k_diffuse_ values (20,000 nm^2^/s, 40,000 nm^2^/s, 100,000 nm^2^/s). (E) Effect of varying k_detach_ while setting k_diffuse_ = 40,000 nm^2^/s. Force versus overlap length are shown for a range of k_detach_ values (0.002 s^−1^, 0.005 s^−1^, 0.01 s^−1^, and 0.05 s^−1^). (F) Numerical simulations were performed using parameter values k_detach_ = 0.002 s^−1^, k_diffuse_ = 40,000 nm^2^/s, and k_end_ = 75. Averaged force values from N=20 independent simulations were calculated and plotted as a function of PRC1 crosslinking molecules. (G) Slopes of data from (F) are plotted as a function of sliding velocity. (H) A schematic model describing how resistive forces increase with both sliding velocity and crosslinker number.

We first sought to explore the model’s parameter space to determine which properties of PRC1 best describe our data. We first modulated k_end_ over several orders of magnitude and express this variation as ratio between k_end_ and k_diffuse_. For small values of k_end_ (e.g. k_end_ = k_diffuse_/500), molecules of PRC1 remained highly clustered and localized near microtubule tips, which acted as reflective barriers against diffusion, producing extremely large forces as overlap length decreased to zero. In contrast, larger values of k_end_ (e.g. k_end_ = k_diffuse_/10) resulted in molecules rapidly diffusing off microtubule ends into solution, producing resistive forces that decreased to zero with decreasing overlap. (Figure 5B).

An intermediate value of ∼50-100 allowed us to reproduce the relatively flat force versus overlap relationship (Figure 5C). We next varied both the diffusion and detachment rates, k_diffuse_(F) and k_detach_(F) respectively. For fast diffusion and detachment, we observed that the forces generated were low and decreased towards zero pN (Figure 5D). As the diffusion rate and detachment rate values were increased, we found that resistive forces increased and the slope of the force relative to overlap length became shallower (Figure 5E). From these simulations, we conclude that the magnitude of resistive force and dependence on parameters such as number of crosslinking molecules and overlap length is highly sensitive to the rate at which molecules will diffuse both within the overlap and off of the microtubule ends, as well as how likely they are to detach from the lattice under load.

Once an optimal range of parameters had been identified, we next sought to determine whether this simple set of conditions was sufficient to recapitulate the velocity- and crosslinker number-dependent resistive force generation behavior that we had observed experimentally. We performed simulations over a range of relative microtubule sliding velocities and determined the magnitude of force as a function of engaged crosslinkers. We found a strong linear relationship between these parameters at all velocities measured (Figure 5F). Slopes from linear regressions on these relationships revealed that the magnitude of resistive force per PRC1 molecule increased linearly with the applied sliding rate (Figure 5G). Together, these results reveal that a simple set of rates that include diffusion on the microtubule lattice, detachment, and diffusion off of microtubule ends are sufficient to describe how resistive forces within sliding bundles crosslinked by PRC1 are generated and depend on sliding velocity and total PRC1 concentration, and correlate well with our experimental measurements.

## Discussion

Our data reveal that ensembles of PRC1 molecules can produce resistive forces against relative microtubule sliding. The magnitude of these forces scales with both the rate of sliding and the total number of engaged crosslinks, but not crosslinker density or overlap length (Figure 5H). The linearly proportional dependence of resistive forces on the sliding velocity suggests that PRC1 ensembles behave as viscous crosslinkers. Upon cessation of sliding the resistive force relaxes, while resumption of sliding results in resistive forces whose magnitude is higher than during the initial sliding event. Finally, we present a simple quantitative model that recapitulates key features of our data by introducing a parameter that allows for microtubule ends to act as partially reflective barriers against PRC1 diffusion.

Based on our results, we suggest that PRC1 ensembles act as a dashpot against microtubule separation. Dashpots are mechanical devices that produce viscous resistance to motions and whose force magnitude is proportional to velocity. We propose that this property may allow sliding microtubules within the spindle midzone to differentially resist motor proteins whose natural stepping rates can span a wide range. For example, recent evidence suggests that microtubule minus-ends can recruit dynein motor proteins via the non-motor MAP NuMA (Charlebois et al., 2011; Elting et al., 2014, 2017). Dynein has been shown to step at rates ∼1 micron/second in vitro; if it were to step at this rate while pulling on microtubule minus-ends, the resistive forces experienced by the motors from the PRC1-mediated overlap would be large. In contrast, motor proteins that localize within overlapping bundles within the spindle midzone, such as kinesin-5 or kinesin-4, step at rates of 25-300 nm/s. At these velocities, PRC1 ensembles would produce a lower resistance. Indeed, experiments performed with mixtures of PRC1 and kinesin-5 suggest that PRC1 crosslinks do not substantially slow relative filament sliding (Subramanian et al., 2010).

Recent reports suggest that the yeast MAP65, Ase1, is capable of generating significant forces via an entropic mechanism as molecules are condensed into small overlaps (Lansky et al., 2015). We find that PRC1 does not exhibit this behavior. What might account for the difference between these two similar types of crosslinking molecule? First, the lifetime of Ase1 within overlaps is nearly at least 10-fold longer than PRC1 molecules (Pamula et al., 2019; Schuyler et al., 2003). Second, our computational models recapitulate observed sliding mechanics when the end-diffusion parameter is 50-100 times smaller in magnitude than the lattice diffusion parameter. Decreasing the end-diffusion constant shifts simulated crosslinker behavior to a pattern more similar to Ase1, where crosslinkers can elastically return a partially slid bundle back to its original position via entropic pressure. This suggests that the end-diffusion parameter is the basis of the mechanistic difference between PRC1 and Ase1 bundles. While the authors analogize Ase1’s behavior within overlaps to that of a one-dimensional gas compressed via a piston-like mechanism (Lansky et al., 2015), we propose that PRC1 behaves as if it were compressed by a ‘leaky’ piston, where molecules can be scooped by the reflective barrier of microtubule plus-ends within the shrinking overlap to generate higher packing densities, but a significant fraction of PRC1 molecules do diffuse off the microtubule ends, thereby reducing the contribution from entropic pressure.

We observe that pausing relative microtubule sliding results in a reduction of resistive forces. Upon resumption of sliding, the rate at which force builds up as well as the magnitude of resistive force during the sliding event both increase. We suggest that the dissipation of force during the pause is due to the PRC1 molecules undergoing diffusion within the overlap to relieve the tension built up across individual molecules. These molecules diffuse towards microtubule crosslinking configurations wherein the relative distance between binding sites is minimized, thus minimizing the force contribution. Why might resumption of sliding after this molecular rearrangement result in higher forces? We propose that within the overlaps and given sufficient time, a subset of PRC1 molecules might diffuse into adjacent sites and form higher order complexes. Evidence for multimerization of crosslinkers has been shown for Ase1 diffusing within microtubule overlaps (Kapitein et al., 2008). In these experiments, diffusive Ase1 molecules will undergo a random walk within overlap until colliding with nearby Ase1 molecules, at which point a brighter cluster with a slower diffusion rate is observed, suggesting two or more molecules have interacted to form a type of complex. We propose that a similar mechanism might be at play in our assays, where multiple PRC1 molecules within non-moving bundles could interact to form higher order assemblies which would likely contribute to the observed higher resistive forces. Recent high-resolution analyses of midzone formation reveal that shrinking microtubule overlaps appear to slow down throughout anaphase, nearly stopping their motion even while spindle poles still separate (Pamula et al., 2019). We hypothesize that were higher-order PRC1 clusters to form during this slow-down or pausing phase, it would lead to an increased resistive load against further microtubule sliding, thereby contributing to the stabilization and maintenance of a robust spindle midzone bundle.

Together our results elucidate mechanisms by which ensembles of PRC1 molecules can regulate the speed of microtubule motions within dividing cells. It will be interesting to see if other non-motor proteins that have been shown to bundle and cluster spindle or kinetochore microtubules, such as NuMA or HURP, produce similar viscous forces against microtubule sliding, or whether an alternate mechanism is employed. It will similarly be interesting to determine whether similar mechanical dashpots are utilized within other cellular processes, such as during remodeling of the actin cytoskeleton.

## Acknowledgements

We thank Dr. Susan Gilbert for the kind gift of the kinesin-1 K439 plasmid and Dr. Blanca Barquera for assistance with construct design. We wish to thank Dr. Brandon Bensel for assistance with the optical trap operation. We thank members of the Gilbert and Bentley labs at RPI, as well as Dr. Radhika Subramanian (MGH) and Dr. Tarun Kapoor (Rockefeller) for helpful discussions, and members of the Forth lab for critical reading of the manuscript. The authors declare no competing financial interests. Author contributions: I.G., A.A. and S.F designed experiments; I.G. and A.A. collected data; M.A. and S.F. designed and performed simulations; I.G. and S.F. wrote manuscript. Funding was provided by RPI School of Science start-up funds to S.F.

---

**Supplementary Figure 1.**
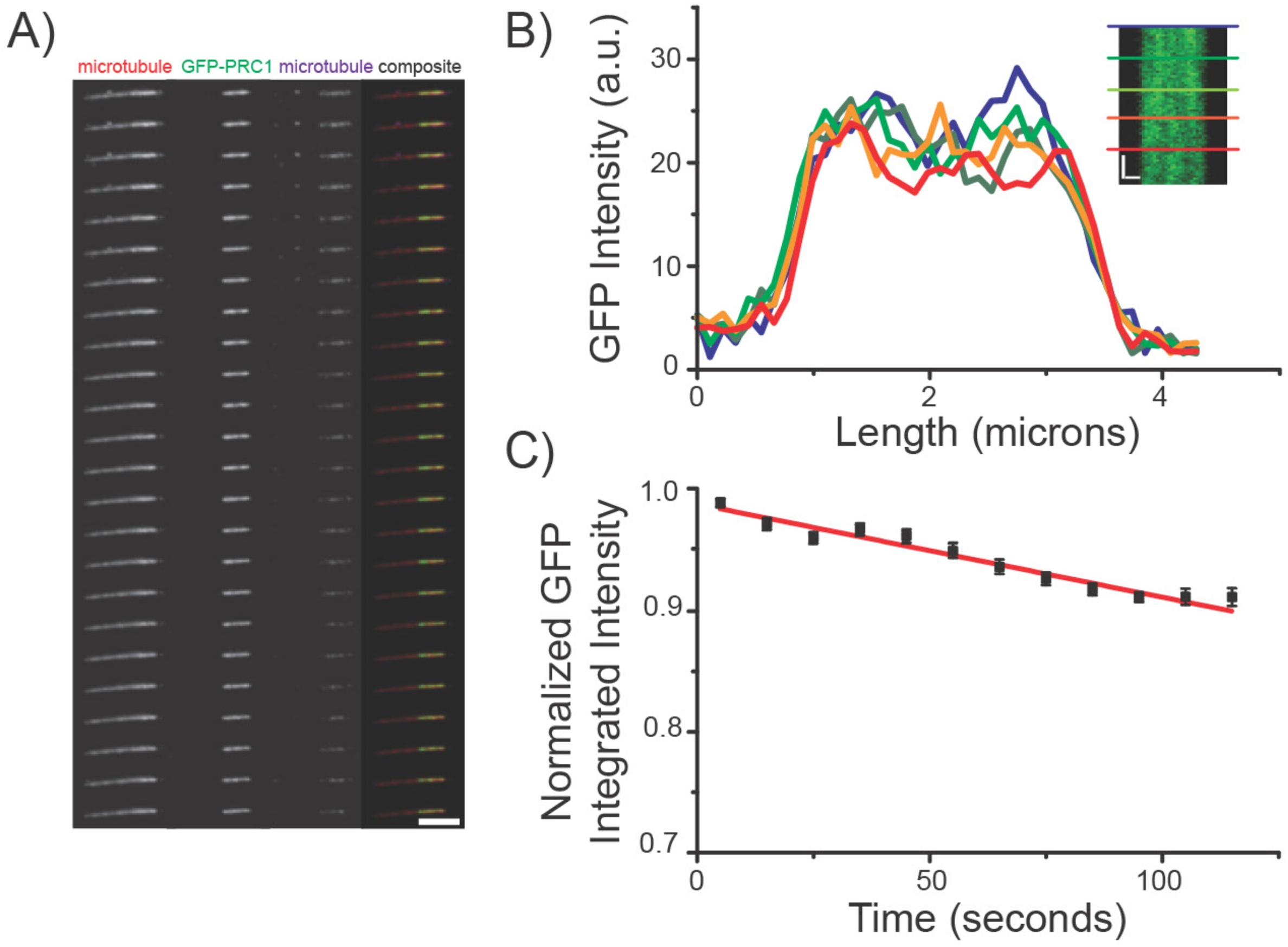
Direct Measurement of GFP-PRC1 Fluorescence Signal Reduction due to Photobleaching. Assay set up similar to description in Figure 1 without kinesin coated bead or stage motion. (A) Time series of fluorescent images acquired via TIRF microscopy. GFP-PRC1 and individual microtubule channels are shown along with composite image showing every 3^rd^ frame (∼5.25 second interval). Scale bar = 2 microns. (B) Linescan analysis from GFP channel. GFP intensities at select time points are plotted against length, with corresponding time points noted on kymograph inset (blue = 0s, green = 26s, yellow = 50s, orange =74s, red = 98s). Kymograph scale bars: vertical = 17s, horizontal = 0.5 microns. (C) GFP signals within overlap regions were integrated and normalized to their initial values and are binned at 10 second intervals. Data from 9 individual measurements were averaged. Error bars = SE.

**Supplementary Figure 2.**
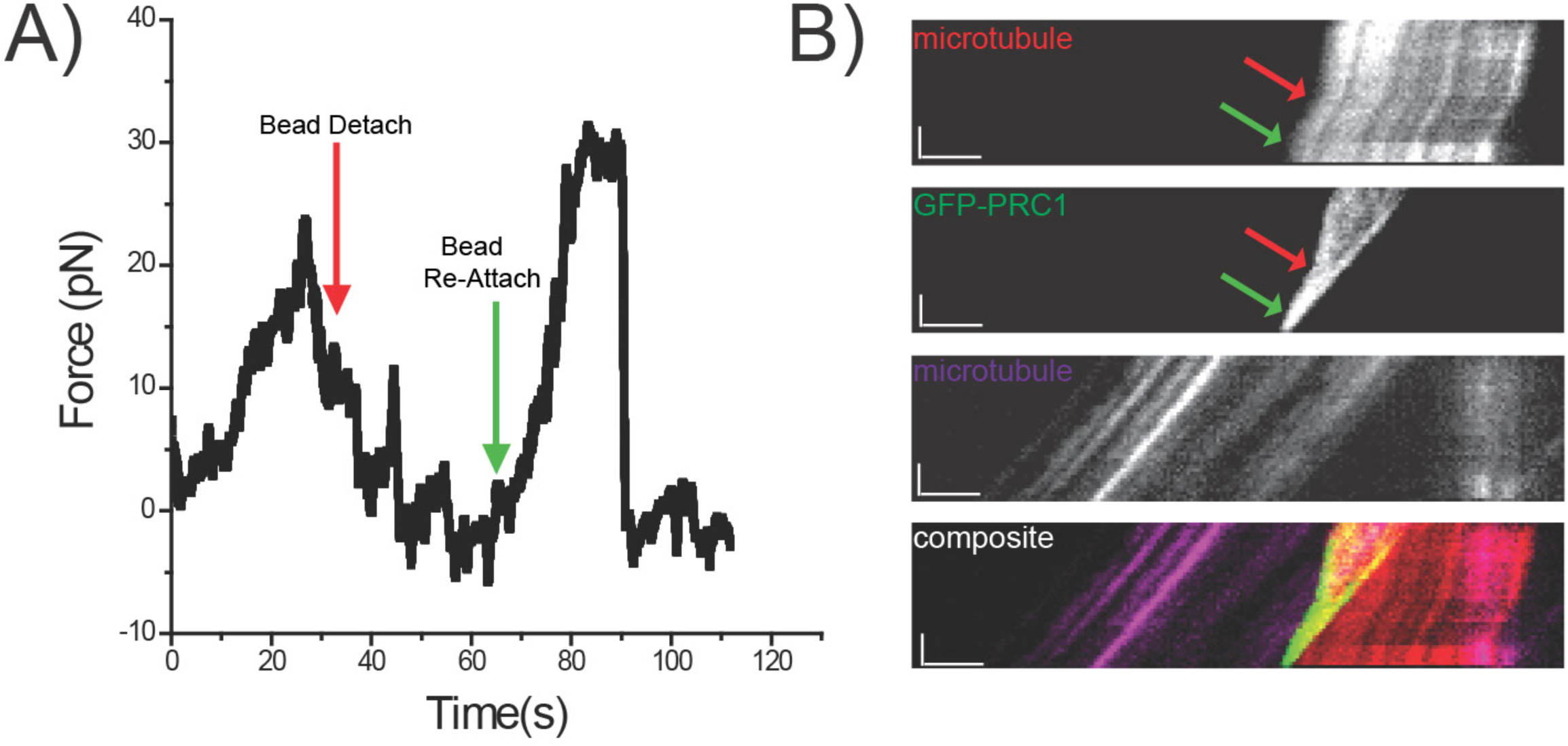
Example of Bead-microtubule detachment and reattachment event. (A) Representative force time series for bead sliding along microtubule assay. Red arrow indicates kinesin coated bead detachment from rhodamine microtubule. Green Arrow indicates kinesin coated bead re-attachment to rhodamine microtubule. (B) Kymographs of corresponding force time series from top to bottom: free rhodamine microtubule, GFP-PRC1, surface-immobilized Hilyte-647 microtubule, and composite of all three channels: rhodamine (red), GFP-PRC1 (green), and Hilyte-647 (purple). Red arrow indicates kinesin coated bead detachment from rhodamine microtubule. Green Arrow indicates kinesin coated bead re-attachment to rhodamine microtubule. Kymograph scale bars: vertical = 19s, horizontal = 2 μm.

**Supplementary Figure 3.**
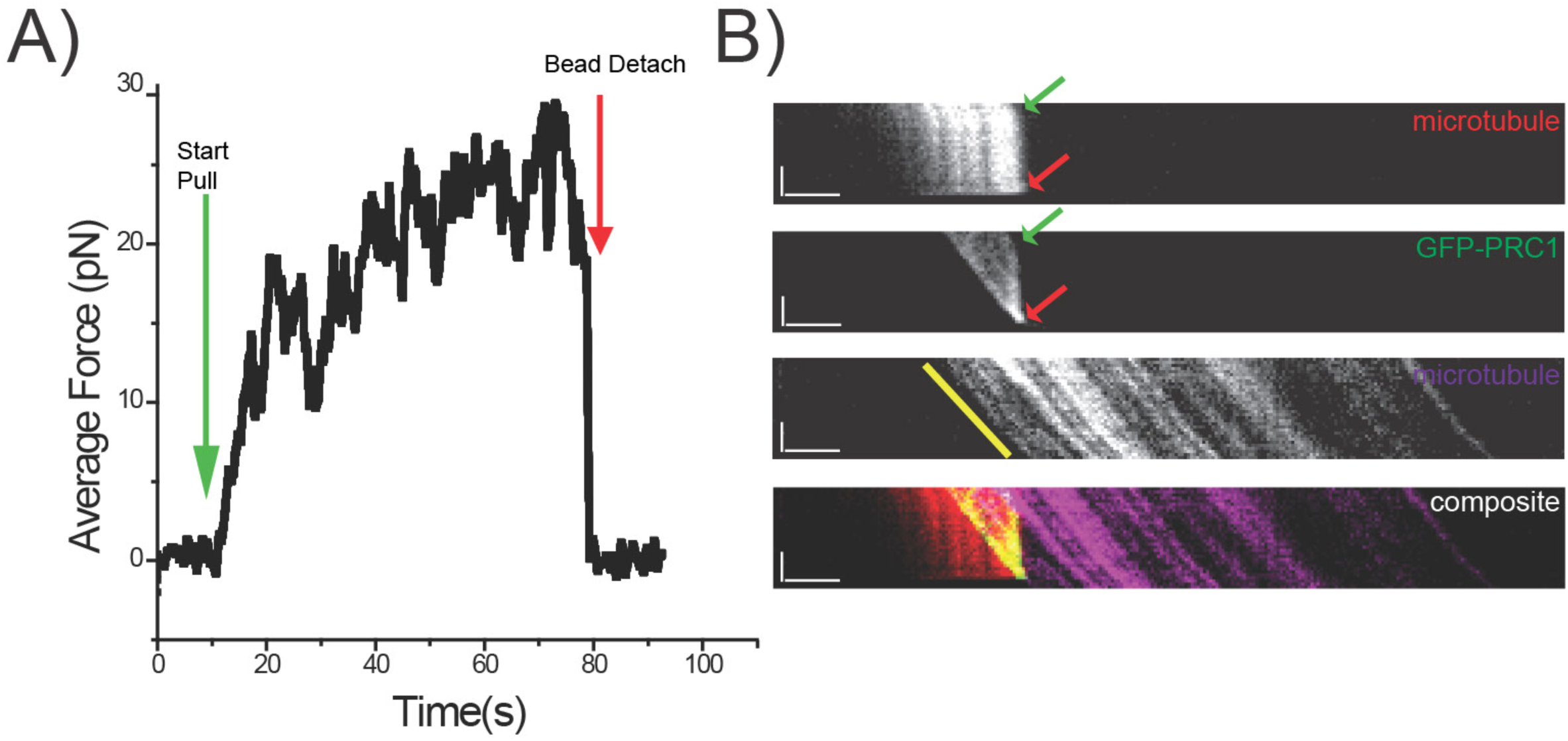
Confirmation of surface microtubule immobilization during sliding event. (A) Representative force time series for bead sliding along microtubule assay. Green arrow indicates start of bundle pull and red arrow indicates bundle separation. (B) Kymographs of corresponding force time series from top to bottom: free rhodamine microtubule, GFP-PRC1, surface-immobilized Hilyte-647 microtubule, and composite of all three channels: rhodamine (red), GFP-PRC1 (green), and Hilyte-647 (purple). Green arrow indicates start of bundle pull and red arrow indicates bundle separation. Yellow line highlights trajectory of surface bound HiLyte647 microtubule and corresponds to a measured velocity of 50nm/s. Kymograph scale bars: vertical = 19s, horizontal = 2 μm.

## Methods

### Protein expression and purification PRC1

We first generated an GFP-tagged PRC1 isoform 2 (aa:1-606) construct by deleting 14aa from near the C-terminus of PRC1 isoform 1 (aa:1-620) using Quikchange mutagenesis (Agilent). The PRC1 isoform 2 gene was then in a pET-DUET plasmid containing an N-terminal histidine tag followed by a Tobacco Etch Virus (TEV) cleavage site and an EGFP sequence inserted in between the TEV cleavage site and the N-terminus of PRC1 isoform 2 with a ‘AAA’ linker sequence just after the eGFP (Subramanian et al., 2010). GFP-PRC1 isoform 2 proteins were expressed via BL21(DE3) Rosetta *Escherichia coli* cells (Novagen). Full length PRC1 was expressed for 4 hours at 18 °C after induction with 0.5mM IPTG. Cells were lysed via sonication in lysis buffer (1 mg/mL lysozyme, 50 mM phosphate (pH 8.0), 10 mM imidazole, and 1% Igepal and HALT protease inhibitor (Pierce), 2 mM TCEP Bond Breaker, 2 mM Benz-HCl, 1 mM PMSF). The lysate was clarified by ultracentrifugation and the supernatant was incubated with Ni-NTA for 1 hour at 4°C (G Biosciences). The resin was then washed with Wash Buffer (50 mM phosphate, (pH 8), 500 mM KCl, 10 mM imidazole, 0.1 % tween, 0.5 mM TCEP, 1 mM PMSF) and the protein was then eluted using Elution Buffer (50 mM phosphate (pH 7.0), 250 mM imidazole, 150 mM KCl, 0.5 mM TCEP). Elute was pooled and concentrated to volume of approximately 1 mL. Next, 1/30 w/w Pro TEV protease (PROMEGA), 1 mM DTT, and 50 μL of Pro TEV 20X buffer were added and the sample incubated in a 30°C water bath for 15 mins before overnight dialysis at 4°C in Gel Filtration Buffer (1X BRB80 (pH 6.8), 150 mM KCl, 10 mM βME). Size exclusion chromatography (Superose-6 increase 10/300 column, GE Healthcare) was then performed in Gel Filtration Buffer with a Shimadzu High Performance Liquid Chromatograph. After collecting peak fractions and concentrating to ∼0.5 mg/mL, sucrose was added to 35% w/v before flash freezing in liquid nitrogen.

### Kinesin-1 K439

Plasmid encoding truncated kinesin-1 construct (K439) with a C-terminally fused EB1 sequence to create stable dimers and a His_8_ tag (Woll et al., 2018) was generously donated by the Dr. Susan Gilbert lab. The protein expression protocol was identical to PRC1, except induction occurred overnight for 18 hours at 16°C. The pellet was resuspended in lysis buffer (10 mM NaPO_4_ (pH 7.2), 300 mM NaCl, 2 mM MgCl_2_, 0.1 mM EGTA, 10 mM PMSF, 1 mM DTT, 0.2 mM ATP, 1 mg/mL lysozyme, 30 mM imidazole, 3 μL benzonase (Novagen) and sonicated. The lysate was clarified by ultracentrifugation and the supernatant incubated with Ni-NTA for 1 hour at 4°C (G Biosciences). The resin was then washed with Wash Buffer (20 mM NaPO_4_ (pH 7.2), 300 mM NaCl, 2 mM MgCl_2_, 0.1 mM EGTA, 0.02 mM ATP, 50 mM Imidazole) and the protein was then eluted using Elution Buffer (20 mM NaPO_4_ (pH 7.2), 300 mM NaCl, 2 mM MgCL_2_, 0.1 mM EGTA, 1 mM DTT, 0.02 mM ATP, 400 mM imidazole). Elute was pooled, concentrated to volume of approximately 1 mL. The protein was then dialyzed overnight at 4°C in Dialysis Buffer (20 mM HEPES (pH 7.2), 300 mM NaCl, 0.1 mM EGTA, 0.1 mM EDTA, 5 mM MgAc, 50 mM KAc, 1 mM DTT. The next day the protein was dialyzed for 1 hour at 4°C in Buffer 1 (20 mM HEPES (pH 7.2), 200 mM NaCl, 0.1 mM EGTA, 0.1 mM EDTA, 5 mM MgAc, 50 mM KAc,1 mM DTT) followed by dialysis for 1 hour at 4°C in Buffer 2 (20 mM HEPES (pH 7.2), 150 mM NaCl, 0.1 mM EGTA, 0.1 mM EDTA, 5 mM MgAc, 50 mM KAc,1 mM DTT) Further purification was performed using size exclusion chromatography (Superose-6 increase, GE Healthcare) using Shimadzu High Performance Liquid Chromatograph. Peak fractions were pooled and dialyze for 3 hours at 4°C of Buffer 3 (20 mM HEPES (pH 7.2), 100 mM NaCl, 0.1 mM EGTA, 0.1 mM EDTA, 5 mM MgAc, 50 mM KAc,1 mM DTT, 5% Sucrose). Protein was flash frozen in liquid nitrogen.

### Microtubule preparation

Microtubule tubulin reagents were purchased from Cytoskeleton, Inc. Microtubules for surface-immobilization were generated via mixture of HiLyte 647 tubulin (TL670M), biotinylated tubulin (T333P), and unmodified tubulin (T240) at a ratio of 1:1:20 along with 1mM GMPCPP. Microtubules were polymerized at 37°C for 1 hour before clarification and stabilization in 30uM Taxol following published protocols (Shimamoto 2015). ‘Free’ microtubules for attachment to optically trapped beads were generated via mixture of rhodamine tubulin (TL590M) and unmodified tubulin at a ratio of 1:20 along with 1mM GMPCPP, and were polymerized, clarified, and stabilized in 30uM Taxol following similar protocols.

### Coating Beads with K439

One-micron diameter streptavidin-coated polystyrene beads (Polysciences Inc., #24162-1) were coated with biotin-conjugated His_6_ antibody (Invitrogen) followed by washing and storage in 2mg/mL alpha-Casein solution made in BRB80 (80 mM K-PIPES, 1mM MgCl_2_, 1mM EGTA, 2mM DTT (pH 6.8)). K439 samples were diluted into BRB80 mixed at 1:1 ratio with coated polystyrene beads. Samples were placed on a rotor for 30 minutes at 4°C followed by sonication for 10 minutes.

### Flow Chamber Construction

The flow chamber design and assay preparation were modified from a previously described protocol (Shimamoto et al., 2015). Anti-parallel microtubule bundles were constructed using passivated glass coverslips coated with SVA-PEG at a ratio of 50 PEG:1 biotin-PEG. All reagents were prepared with BRB80 buffer. Following each reagent flow-in and incubation, a flush with ∼3 chamber volumes of BRB80 was performed. Reagents were introduced stepwise with the following order and incubation times: (1) 0.5 mg/mL neutravidin, 2 minutes; (2) 0.5 mg/mL alpha casein surface block, 3 minutes; (3) HiLyte-647 biotinylated microtubules with 0.2mg/mL alpha casein with no additional incubation and immediate flush; (4) 1 nM GFP-PRC1 with 0.2 mg/mL alpha casein, 2 minutes; (5) Rhodamine 561 microtubules with 0.2 mg/mL alpha casein, 5 minutes, and the corresponding chamber flush included 1 mM TCEP bond breaker solution in BRB80; (6) 1 μL of K439 coated beads, 1 mM of TCEP bond breaker, 0.2 mg/mL alpha casein, 70 mM KCl, Oxygen Scavenging System (4.5 mg/ml glucose, 350 U/ml glucose oxidase, 34 U/ml catalase, 1 mM DTT). The chamber was then sealed with clear nail polish prior to experiments.

### Image Acquisition

Microtubule bundles were imaged using three-channel TIRF microscopy using the following laser lines and exposure times: HiLyte-647 microtubules: 640 nm laser (60% power, 200 ms exposure); GFP-tagged PRC1: 488 nm laser (30% power, 100 ms exposure); and rhodamine microtubules: 561 nm laser (30% power, 100 ms exposure). Images were acquired using a Photometric Prime 95B camera controlled with Nikon NIS Elements software at overall acquisition rates of one frame per ∼1.75 seconds. Prior to analysis, images were visually screened to ensure that there were no additional interactions with microtubules from other bundles. Analysis of fluorescent data and generation of intensity linescan data sets were performed using a combination of FIJI (ImageJ) tools and custom-written LabVIEW software.

### Force Data Acquisition

The optical tweezers system was constructed based on a fiber-coupled infrared laser (1064 nm, 10W, IPG Photonics) and a position-sensitive detector (Thorlabs PDQ80A). The laser beam power was controlled using an AOM (AA Optoelectronics MTS80-A3-1064Ac) and expanded using a custom-built telescope (Thorlabs optics). The beam was then passed through a 1:1 telescope and merged into the microscope’s light path using a dichroic IR filter (z900dscp, Chroma) mounted on a secondary turret located above the fluorescence turret. The beam was introduced into the back aperture of the objective and focused to a diffraction-limited spot such that a micron-sized bead could be trapped approximately 500nm above the coverslip surface. A high NA 100× objective (1.49 NA; CFI Plan Apo TIRF) was used for establishing both stable optical trapping and high-resolution fluorescence imaging. Coverslip position was controlled with sub-nanometer precision using a closed-loop three-axis piezo stage (Nano LPS-200, Mad City Labs). To monitor the displacement of the bead held in the optical tweezers, the laser beam passing through the sample plane was collected using an oil-immersion condenser (MEL41410, Nikon) and reflected away from the microscope’s imaging path by a dichroic IR mirror (DMSP805L, Thorlabs). After passing through a long-pass optical filter, the beam was projected onto the quadrant photodiode detector placed conjugate to the back-focal plane of the objective. The signals from each photodiode quadrant were amplified and recorded using an in-house developed LabVIEW program via an AD converter (PCIe-6363, National Instruments). The displacement signal along each coordinate was obtained by calculating the difference of the normalized signals between the adjacent quadrant pairs. Bead position and force data were converted from raw voltage to physical units after standard calibration methods were employed using custom-written LabView software. Absolute force values were determined by introducing a constant offset such that zero-force corresponded to force values recorded immediately upon bundle separation.

### Computational Simulations

To simulate the resistive forces generated by ensembles of diffusive PRC1 crosslinkers between two sliding microtubules, we modified a Monte-Carlo based technique previously employed to model kinesin-5 stepping behavior (Shimamoto et al., 2015). Briefly, individual PRC1 molecules can be defined as two microtubule binding domains connected by a spring-like linker domain that allows for crosslinking of two individual microtubules. For each MT-binding domain a diffusion parameter *k*_*diffuse*_ can be assigned, as well as a rate of detachment *k*_*detach*_, and both parameters can depend on applied force via an Arrhenius-like term.

To describe the behavior of a PRC1 molecule which is crosslinking two microtubules, we consider two independent MT-binding domains connected by a spring-like linker region which stretches as the two heads become spatially separated as they move along the microtubule surface. This parameter *k*_*stiffness*_ is a simple linear spring constant term that coarsely defines an energy penalty for the spatial separation of the MT-binding domains.

#### Distribution of forces across PRC1 molecules with optically trapped microtubule end

The central feature of our simulation is the balance of forces across the PRC1 molecules with the optically trapped microtubule end serving as a linear spring providing resistance to microtubule motion. For a given number of molecules, we require that the forces must balance according to:

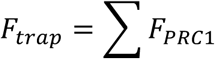

By defining the coordinates of discrete (i.e., 8 nm spacing) positions along the free (‘top’) microtubule and the discrete positions along the immobilized ‘bottom’ microtubule, as well as considering the position of the moving free microtubule end (which is equivalent to the optical trap position, *x*_*trap*_), we can rewrite this relationship as:

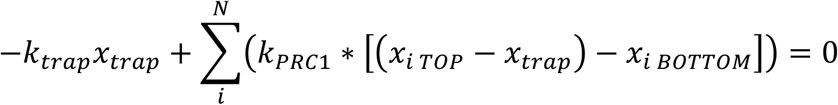

Solving for *x*_*trap*_ yields:

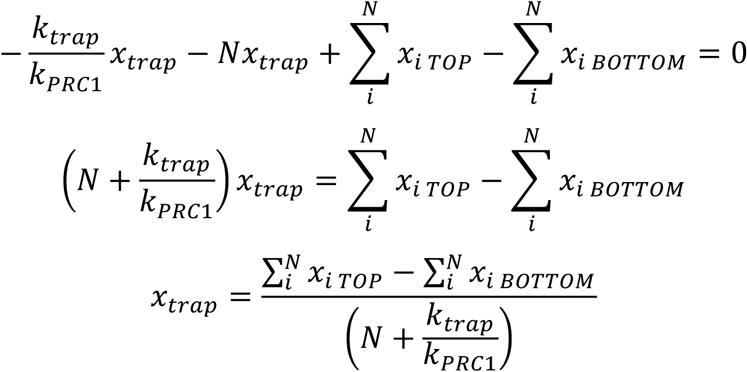

The position of the microtubule end (trap position, *x*_*trap*_) will therefore depend on the relative positions along the microtubules of each PRC1 MT-binding domain, the total number of molecules engaged in crosslinking, and both the trap and PRC1 linker domain stiffness.

#### Monte Carlo Simulations

Start at Time = 0 s with all PRC1 molecules in a fully relaxed state (i.e., with all Force_PRC1_ = 0 pN) and uniformly spaced within the overlap region, as well as the condition that Force_trap_ = 0 pN.

At each subsequent time step, perform the following:

1. For each PRC1 molecule, check the attachment state of both the top and bottom heads. If the head is attached, detach if rand() < *k*_*detach*_(*F*)*Δ*t*. If rand() > Probability, do nothing and calculate attachment state of next molecule.
2. For each diffusive head:
  a. Allow the head to diffuse in the direction of force application if rand()<*k*_*diffuse*_*exp(F*(4 nm)/kT) or in the direction opposite force application if rand()>1-(*k*_*diffuse*_*exp(-F*(4 nm)/kT)).
  b. If adjacent site is occupied by another PRC1 molecule, do not allow head to diffuse onto the site; head remains at current site.
  c. If PRC1 molecule is occupying the last available site on the microtubule lattice, allow it to diffuse off of microtubule end if rand()<*k*_*end*_*exp(F*(4 nm)/kT).
  d. If none of the above conditions are met, allow the head’s position to remain unchanged.
3. Determine the number of PRC1 molecules that have both heads attached, *N*_*engaged*_.
4. Calculate the new position of the trap according to:

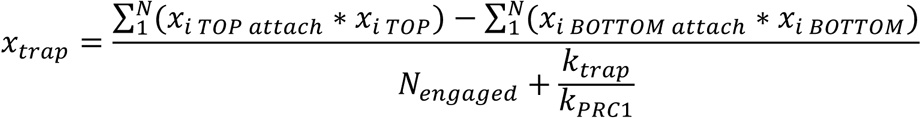
5. Calculate the Force across each attached motor according to:

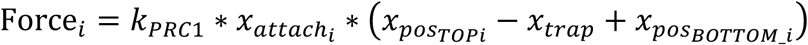
6. Increment the absolute position of the bottom (“surface-immobilized”) microtubule by a distance Rate* Δ*t*, where rate is selected from {25, 50, 100, 200} nm/s. The overlap length is therefore reduced in an approximately linear manner until it reaches zero, at which point no PRC1 molecules are available to crosslink the microtubule pair and the force on the free microtubule as measured by the bead falls to zero.

Repeat steps 1-6 with updated values for Force per PRC1 molecule for desired length of time (typically 30-90 seconds).

